# Characterizing human postprandial metabolic response using multiway data analysis

**DOI:** 10.1101/2023.08.31.555521

**Authors:** Shi Yan, Lu Li, David Horner, Parvaneh Ebrahimi, Bo Chawes, Lars O. Dragsted, Morten A. Rasmussen, Age K. Smilde, Evrim Acar

**Author notes:** M.A.R., A.K.S. and E.A. designed the research; B.C. and M.A.R. curated the data; S.Y. carried out the data analysis; L.L. and E.A. validated the results; D.H., P.E., L.O.D., and M.A.R. interpreted the results; S.Y., A.K.S., and E.A. wrote the paper. The authors declare no competing interest.

## Abstract

Analysis of time-resolved postprandial metabolomics data can enhance our knowledge about human metabolism by providing a better understanding of similarities and differences in postprandial responses of individuals, with the potential to advance precision nutrition and medicine. Traditional data analysis methods focus on clustering methods relying on summaries of data across individuals or use univariate methods analyzing one metabolite at a time. However, they fail to provide a compact summary revealing the underlying patterns, i.e., groups of subjects, clusters of metabolites, and their temporal profiles. In this study, we analyze NMR (Nuclear Magnetic Resonance) spectroscopy measurements of plasma samples collected at multiple time points during a meal challenge test from 299 individuals from the COPSAC_2000_ cohort. We arrange the data as a three-way array: *subjects* by *metabolites* by *time*, and use the CANDECOMP/PARAFAC (CP) tensor factorization model to capture the underlying patterns. We analyze the *fasting state* data to reveal static patterns of subject group differences, and the *fasting state*-corrected postprandial data to reveal dynamic markers of group differences. Our analysis demonstrates that the CP model reveals replicable and biologically meaningful patterns capturing certain metabolite groups and their temporal profiles, and showing differences among males according to their body mass index (BMI). Furthermore, we observe that certain lipoproteins relate to the group difference differently in the fasting vs. dynamic state in males. While similar dynamic patterns are observed in response to the challenge test in males and females, the BMI-related group difference is only observed in males in the dynamic state.

---

**F**ood ingestion triggers many parts of the human metabolism. While the fasting state may reveal certain metabolic differences among individuals, recently *challenge tests* have been used to study also the *postprandial* metabolism of individuals. Analyzing postprandial data may help us understand differences, stratify individuals in terms of their metabolic responses (1, 2) and advance personalized nutrition (3, 4). For example, postprandial hyperlipidemia, the abnormally increased levels of triglyceride-rich lipoproteins after food intake, is a risk factor for cardiovascular diseases (5–7). A postprandial study on type 2 diabetes (T2D) gives insight into the flexibility on the human metabolome (8) and subjects with different body mass index (BMI) values have shown different postprandial responses in terms of amino acids (9, 10) and lipid metabolites (11). Sex differences in insulin sensitivity and metabolome have been observed in the postprandial response (12) (See (13) for a recent review on challenge tests and metabolomic responses reported in the literature.) To take it even further, personalized dietary recommendations based on integrating postprandial data with, e.g., gut microbiome characteristics, may lower the postprandial glucose levels (3).

The state-of-the-art methods to analyze challenge test data can be divided into *supervised* and *unsupervised* methods. In many cases, there is a study design underlying a challenge test with different groups (12, 14, 15). For instance, there can be groups as a result of baseline characteristics, e.g., normal vs. hyper-triglycerides (14), or healthy vs. diseased or pathological conditions (12). The data in such studies is labelled, and often analyzed using *supervised* methods. If there is no a priori grouping nor an underlying treatment, then such data is unlabelled, and analyzed using *unsupervised* methods. An example is cohort data; the topic of this paper.

The predominant way to analyze *labelled* data is to use univariate statistical methods in which the metabolites are analyzed separately. Depending on the exact background of the study design, such methods can be *t*-tests (14, 16) or analysis of variance (ANOVA) models (4, 12, 14, 17, 18) to study group differences. More advanced methods consider explicitly the temporal aspect of the data and/or unbalanced designs, e.g., by using linear mixed models (19), extensions of the analysis of variance-simultaneous component analysis (ASCA) (20). A shortcut for analyzing the temporal behavior is sometimes used by extracting features from the dynamic profiles such as the area under the curve (AUC) (11, 21, 22).

## Significance Statement

Analysis of postprandial metabolomics data can improve our understanding of variations among individuals in terms of metabolic responses, and holds the promise to reveal static and dynamic markers of metabolic diseases. Time-resolved postprandial metabolomics data is in the form of a three-way array: *subjects* by *metabolites* by *time*. Traditional data analysis approaches fail to preserve the multiway structure of such data. We use tensor factorizations to analyze three-way postprandial metabolomics data and reveal the underlying patterns. Our analysis demonstrates that tensor factorizations reveal replicable and biologically meaningful patterns capturing metabolite groups, their temporal profiles, and group differences (among males according to their body mass index). We discuss observed differences in static vs. dynamic markers as well as males vs. females.

Although all these methods are powerful, they rely on labelled data, and the univariate ones do not take into account the correlation between metabolites.

The predominant *unsupervised* analysis of challenge test data is by using multivariate data analysis methods such as principal component analysis (PCA) or cluster analysis. PCA has been used to analyze measurements from all time points and subjects by arranging the data as *subjects-time points* by *metabolites* (23) or one time point at a time (24). Clustering is performed on time profiles of all metabolites (e.g., averaged across subjects) to create groups of metabolites showing a similar time course (2, 22). These multivariate methods work by considering only summaries of the data and/or do not utilize the fact that challenge test data has a more complex and temporal structure.

Postprandial metabolomics data collected at multiple time points from multiple subjects is inherently three-way, and can be arranged as a *subjects* by *metabolites* by *time points* array (see Fig. 1). Multiway data analysis methods (also known as tensor factorizations) (25–27) are effective tools for extracting the underlying patterns from such multiway data (also referred to as higher-order tensors), and have been successfully used in neuroimaging data analysis (28, 29), electronic health records-based phenotyping (30, 31) and chemometrics (32, 33).

**Fig. 1.**
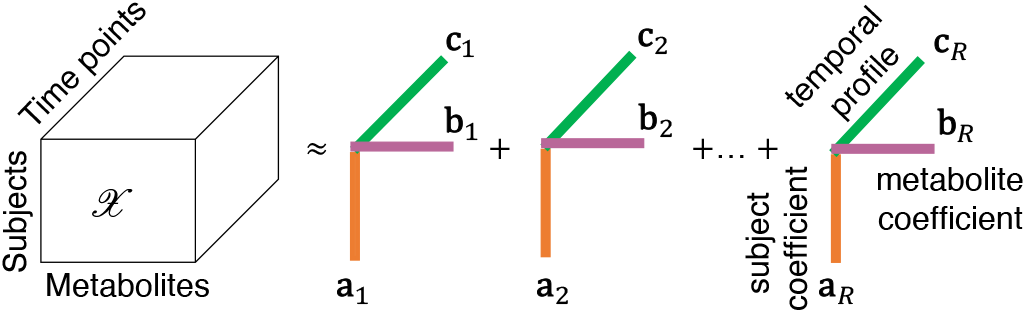
An *R*-component CP model of a third-order tensor with modes: *subjects, metabolites*, and *time points*.

With the goal of stratifying subjects with no prior labels and understanding differences among subjects based on their metabolic responses to a meal, in this paper, using unsupervised methods, we analyze measurements of plasma samples collected during a meal challenge test from the COPSAC_2000_ cohort (Copenhagen Prospective Studies on Asthma in Childhood) (34). We arrange the metabolomics measurements at several time points from a group of participants as a *subjects* by *metabolites* by *time points* tensor, and use the CANDECOMP/PARAFAC(CP) tensor factorization model (35, 36) to reveal the underlying patterns in the data. The CP model summarizes the data by extracting the main patterns of variation in the *subjects, metabolites* and *time* modes simultaneously. Extracted patterns are unique under mild conditions (25, 37) and this facilitates interpretation. While CP-based tensor methods have been previously used in longitudinal data analysis, e.g., analysis of gut microbiome (38), urine metabolomics (39), and simulated dynamic metabolomics data (40), its application in postprandial metabolomics data analysis has been limited. We recently simulated postprandial metabolomics data using a human whole-body metabolic model (41), and demonstrated that analysis of the T0-corrected data (i.e., the postprandial data corrected by subtracting the fasting state data, similar to the method of analysis of changes (42)) using a CP model together with the analysis of the fasting state data provides a comprehensive picture of the underlying metabolic mechanisms.

In this paper, building onto previous work on simulations (41), we carry out a comprehensive analysis of the time-resolved postprandial metabolomics data collected from the COPSAC_2000_ cohort, and analyze the T0-corrected data using a CP model and the fasting state data using PCA. We demonstrate that the CP model reveals biologically meaningful and replicable patterns (i.e., replicable across subsets of subjects), and use the replicability of the patterns as a selection criterion for the number of components in the CP model.

## 1. Data description

The COPSAC_2000_ cohort contains 411 healthy subjects with mothers with a history of asthma. 299 subjects, aged 18 years, attended the food challenge test using a standardized mixed meal (43) containing 60 g palm olein, 75 g glucose, and 20 g dairy protein in a total volume of 400 ml water-based drink. The size of the drink was made proportional to the daily recommended caloric intake (function of sex, age and height). Blood samples were collected from the participants at eight time points after an overnight fasting, i.e., at the fasting state, and 0.25, 0.5, 1, 1.5, 2, 2.5, and 4 hours after the meal intake. The samples were put on ice, and within 4 hours split into EDTA plasma before being stored at -80 ^*°*^C. The study was conducted in accordance with the Declaration of Helsinki and was approved by the Copenhagen Ethics Committee (KF 01-289/96 and H-16039498) and the Danish Data Protection Agency (2008-41-1754).

Plasma samples were measured using NMR (nuclear magnetic resonance) spectroscopy through Nightingale Blood Biomarker Analysis, which provides 250 features^∗^. These features consist of lipoproteins, apolipoproteins, aminoacids, fatty acids, glycolysis-related metabolites, ketone bodies and an inflammation marker. Lipoproteins consist of four main lipoprotein classes: HDL (high density lipoprotein), IDL (intermediate density lipoprotein), LDL (low density lipoprotein), and VLDL (very low density lipoprotein), and subclasses divided according to particle sizes: XXL (extremely large), XL (very large), L (large), M (medium), S (small) and XS (very small). Cholesterol (C), cholesterol ester (CE), free cholesterol (FC), triglyceride (TG), phospholipid (PL) levels of these classes and subclasses are among the list of metabolites.

In addition, we include measurements of insulin and C-peptide. The total number of features is 161 (see supplementary material S1). Seven subjects are removed before the analysis since two subjects have a large amount of missing data, four subjects have extremely high levels of acetate, and one subject is detected as an outlier for the CP model (for outlier detection, see (32)). In total, there are 140 male and 152 female subjects included in the analysis.

The T0-corrected metabolomics data, i.e., the postprandial data corrected by subtracting the fasting state data, is arranged as a third-order tensor *𝔛* of size *I* × *J* × *K*, where *I, J, K* represents the number of subjects (140 males or 152 females), the number of metabolites (161 features), and the number of time points (7 time points, i.e., fasting state data is subtracted from other time points and excluded in T0-corrected data), respectively. The fasting state data is arranged as a matrix of size *I* × *J*. See Table 1 for the data sets considered in this paper.

**Table 1.** Data sets constructed from the postprandial metabolomics measurements and the corresponding analysis methods.

|  | Subjects | # Time points | Data Size | Analysis method |
| --- | --- | --- | --- | --- |
| Section 3.1 | males | 7 | 140 × 161 × 7 | CP |
| Section 3.2 | males | 1 | 140 × 161 | PCA |
| Section 3.3 | females | 7 | 152 × 161 × 7 | CP |
| Section 3.4 | females | 1 | 152 × 161 | PCA |

Different body composition, and insulin resistance measures were also obtained from the subjects. Weight, height, and waist circumference were directly measured, and consequently BMI and waist/height ratio were calculated. Body composition was estimated by bioelectrical impedance using a Tanita scale (TANITA MC-780MA), to obtain body muscle mass and fat mass, and subsequently, body fat percentage, muscle to fat ratio, fat mass index (fat mass/height^2^ (kg/m^2^)), and fat free mass index ((muscle mass/2.2) * 2.20462/height^2^ (kg/m^2^)). Insulin resistance was assessed using HOMA-IR (Homeostatic Model Assessment for Insulin Resistance), by the formula: fasting insulin (microU/L) *×* fasting glucose (nmol/L) */* 22.5.

## 2. Methods

### A. CP model

The CP model (35, 36, 44) approximates a higher-order tensor as the sum of a minimum number of rank-one tensors as in Fig. 1. An *R*-component CP model of a third-order tensor *𝔛* ∈ ℝ^*I×J×K*^ with modes: *subjects, metabolites*, and *time points* represents the data as follows:

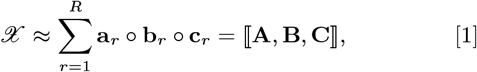

where denotes the vector outer product; **a**_*r*_, **b**_*r*_, **c**_*r*_ corresponds to the *r*th column of factor matrices **A** ∈ ℝ^*I×R*^, **B** ∈ ℝ^*J×R*^, and **C** ∈ ℝ^*K×R*^, respectively. Each component (**a**_*r*_, **b**_*r*_, **c**_*r*_) may reveal subject groups (i.e., subjects with similar coefficients in **a**_*r*_), groups of metabolites related to those subject groups (i.e., metabolites with large coefficients in **b**_*r*_) following a specific temporal profile (given by **c**_*r*_). The CP model is unique up to permutation and scaling ambiguities. Permutation ambiguity indicates that the order of rank-one components is arbitrary while the scaling ambiguity in the model corresponds to arbitrarily scaling each vector in component *r*, i.e., (**a**_*r*_, **b**_*r*_, **c**_*r*_), as long as the product of the norms of the vectors stays the same. Except for these ambiguities, extracted CP patterns are unique and this facilitates the interpretability of the model.

The uniqueness of the CP model without additional constraints on the extracted patterns is an advantage compared to other types of tensor factorizations. For instance, a tensor factorization method called higher-order singular value decomposition (HOSVD) with orthogonality constraints on all factor matrices has recently been used to analyze metabolomics measurements collected at multiple time points before and after an oral glucose challenge test from twenty subjects (45). However, orthogonality constraints in *subjects, metabolites* and/or *time* modes are not realistic, and only needed for the uniqueness of the model. Therefore, the interpretability of such approaches is limited.

The *model fit* (also often referred to as *explained variance*) shows how well the CP model describes the data and is defined as follows:

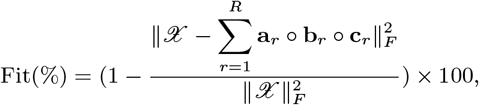

Where ∥ · ∥ _*F*_ denotes the Frobenius norm. If the model fully explains the data, the fit is 100%; otherwise, the unexplained part remains in the residuals.

When reporting the results, we normalize each vector, **a**_*r*_, **b**_*r*_, **c**_*r*_, by its norm due to the scaling ambiguity.

#### A.1. Model Selection

Choosing the right number of components, i.e., *R*, is crucial to extract the underlying patterns accurately; however, the selection of *R* in a CP model is a challenging task (25, 46). Here, we choose *R* based on the *replicability* of the extracted components, where *replicability* is defined as the ability to extract similar patterns from independent subsamples of the dataset (47), and is an extension of split-half analysis (48).

The replicability of an *R*-component CP model is assessed as follows:

Step 1. Randomly split all subjects into ten parts (if there is a class information of potential interest, randomly split the subjects such that each part has the same class proportions as in the original data)

Step 2. Leave out one part at a time, and form a subset using the remaining data, i.e., in total, ten subsets,

Step 3. Fit an *R*-component CP model to each subset,

Step 4. Calculate the similarity of factors between every pair of CP models (in the *metabolites* and *time points* modes), i.e., 45 similarity scores,

Step 5. Repeat Step 1-4 ten times using different random splitting of subjects in Step 1.

The similarity of the factors in *metabolites* and *time points* modes from two CP models ⟦**A, B, C**⟧ and 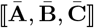 is measured using the factor match score (F MS) d[efined as follows:

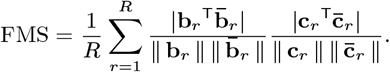

An *R*-component CP model is considered to be replicable if 95% of FMS values (of 450 FMS values computed at the end of Step 5) is higher than 0.9. We choose the highest number of components that produces a replicable CP model in order to explain the data as much as possible (note that the model fit increases as the number of components increases).

### B. Experimental setting

Before the analysis, the data is preprocessed by first centering across the *subjects* mode and then scaling within the *metabolites* mode (49). When scaling, each slice in the *metabolites* mode is divided by the root mean squared value of non-missing entries.

All experiments are performed in MATLAB (2020b). For fitting the CP model, we use `cp-wopt` (50) from the Tensor Toolbox (version 3.1) (51) using the nonlinear conjugate gradient algorithm from the Poblano Toolbox (52). Multiple random initializations are used when fitting the model to avoid local minima. This is also the case when assessing replicability, i.e., multiple random initializations are used when fitting a CP model, and the initialization returning the minimum function value is considered. For PCA, we use the MATLAB function `svd` after estimating the missing values using weighted optimization (50).

## 3. Results

### 3.1. The CP model of T0-corrected metabolomics data from males reveals a BMI-related group difference

We analyze the T0-corrected data from males using a 2-component CP model. The number of components is selected based on the replicability of extracted patterns as shown in Fig. 2. The model fit is 44%.

**Fig. 2.**
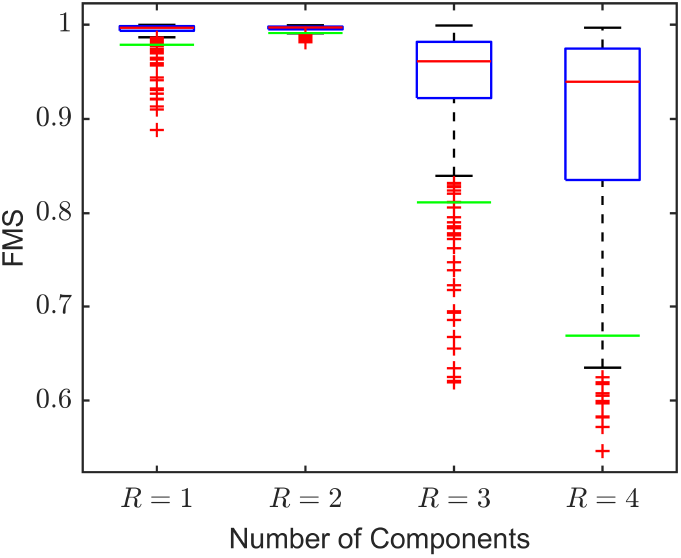
Replicability of the CP model of the T0-corrected data from males for different number of components *R*. Green lines show that 95% of the FMS values are above those lines. Models are replicable for *R* = 1 and *R* = 2. We choose the replicable model with the highest number of components (i.e., *R* = 2).

Fig. 3 shows the factors (**a**_*r*_, **b**_*r*_, **c**_*r*_) extracted by the 2-component CP model. To see whether the model reveals any underlying group structure among subjects, after the analysis, we consider additional information available about the participants. In the *subjects* mode (i.e., **a**_1_ and **a**_2_) in Fig. 3, subjects are colored according to BMI groups, i.e., *lower* vs. *higher* BMI groups. The lower BMI group contains subjects with BMI less than 25.0 (i.e., underweight and normal weight subjects) while the higher BMI group contains subjects with BMI greater than or equal to 25.0 (i.e., overweight and obese subjects). In the *metabolites* mode (i.e., **b**_1_ and **b**_2_), metabolites are colored according to lipoprotein subclasses. Here, the **Rest** group contains features other than the ones in the lipoprotein group while the **Total** group consists of total concentrations of certain metabolites, e.g., Total-C, Total-TG. See supplementary material S1 for details. In the *time* mode (i.e., **c**_1_ and **c**_2_), the model reveals the temporal patterns.

**Fig. 3.**
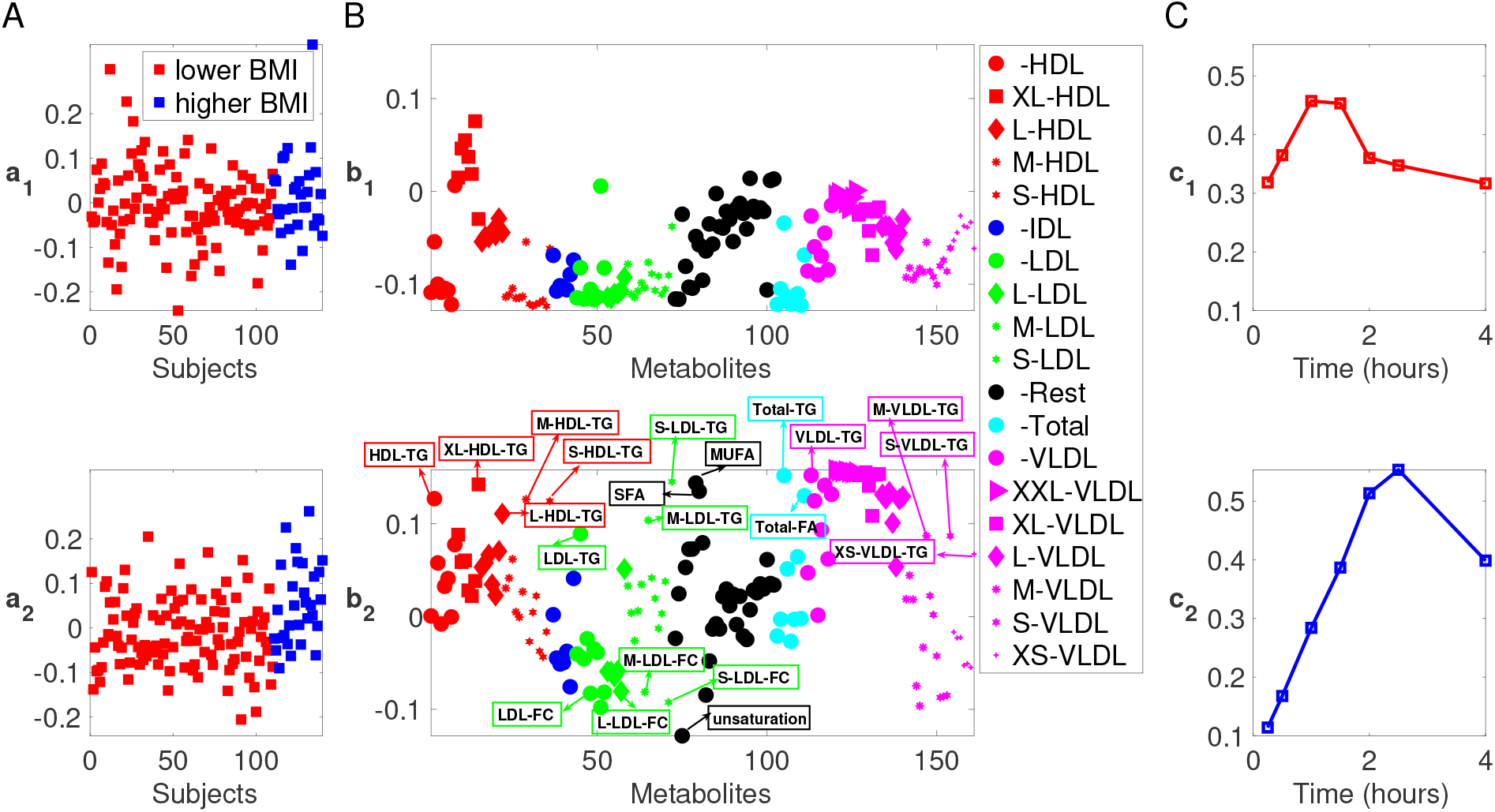
2-component CP model of the T0-corrected metabolomics data from males. (A) *Subjects* mode (i.e., **a**_1_ and **a**_2_), where subjects are colored according to BMI defined as lower BMI: BMI*<* 25 and higher BMI: BMI ≥ 25, (B) *Metabolites* mode (i.e., **b**_1_ and **b**_2_), where metabolites are colored according to lipoprotein classes. The size of the marker for each metabolite is adjusted according to lipoprotein subclasses as indicated in the legend. We show the names of the metabolites with the highest coefficients for the component of interest, i.e., **b**_2_. The Rest group contains features other than the ones in the lipoprotein group, the Total group corresponds to total concentrations of certain metabolites, e.g., Total-C, Total-TG. See supplementary material S1 for details, (C) *Time* mode (i.e., **c**_1_ and **c**_2_).

In this model, the second CP component, i.e., (**a**_2_, **b**_2_, **c**_2_), is of particular interest since it reveals a statistically significant group difference (*p*-value = 6 × 10^*−*4^ using a two-sample *t*-test based on **a**_2_) in terms of BMI (see Fig. 4). In the *metabolites* mode, i.e., **b**_2_ in Fig. 3B, we observe that the subject group difference is due to different behaviour of certain metabolites, i.e., metabolites with high coefficients in terms of absolute value. In particular, we observe that TG-related metabolites, VLDLs (XXL/XL/L), monounsaturated fatty acids (MUFA), saturated fatty acids (SFA), and total fatty acids (Total-FA) have high positive coefficients (indicating that changes in these metabolites positively relate to higher BMI) while several LDL-FCs, VLDLs (M/S), and unsaturation degree have high negative coefficients (indicating that changes in these metabolites negatively relate to higher BMI). Unsaturation degree is the level of unsaturation of fatty acids within a sample, high levels of which indicating that the fatty acid content is likely to be polyunsaturated rather than saturated for a specific sample. In the *time* mode, **c**_2_ increases gradually until 2.5 hours and decreases afterwards showing the temporal profile of the metabolic response modelled by this component.

**Fig. 4.**
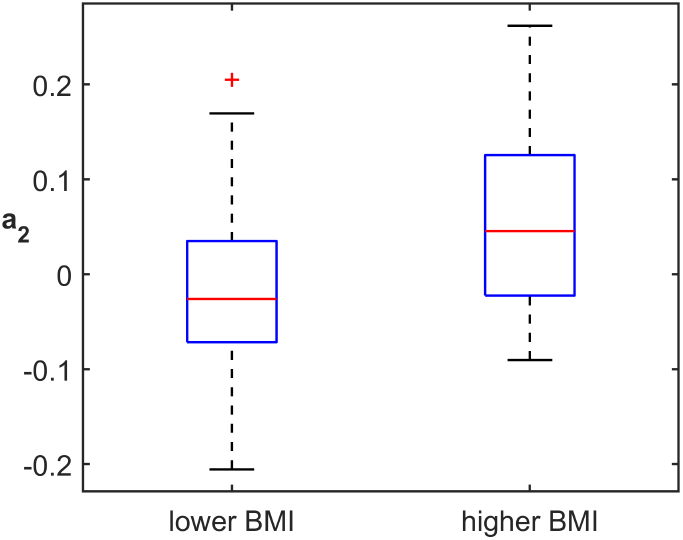
Boxplots corresponding to different BMI groups. The boxplots are plotted based on component 2 in the *subjects* mode, i.e., **a**_2_, extracted by the 2-component CP model of the T0-corrected data from males.

Note that while we mainly discuss subject group differences in terms of BMI values, this component also has significant correlation with several other variables. Fig. 5 shows the correlation of the factor vector in the *subjects* mode for the second component, i.e., **a**_2_, with various variables of interest, by sex. High positive and negative correlations indicate that the second component is related to not only BMI but also a phenotype defined by these closely related variables.

**Fig. 5.**
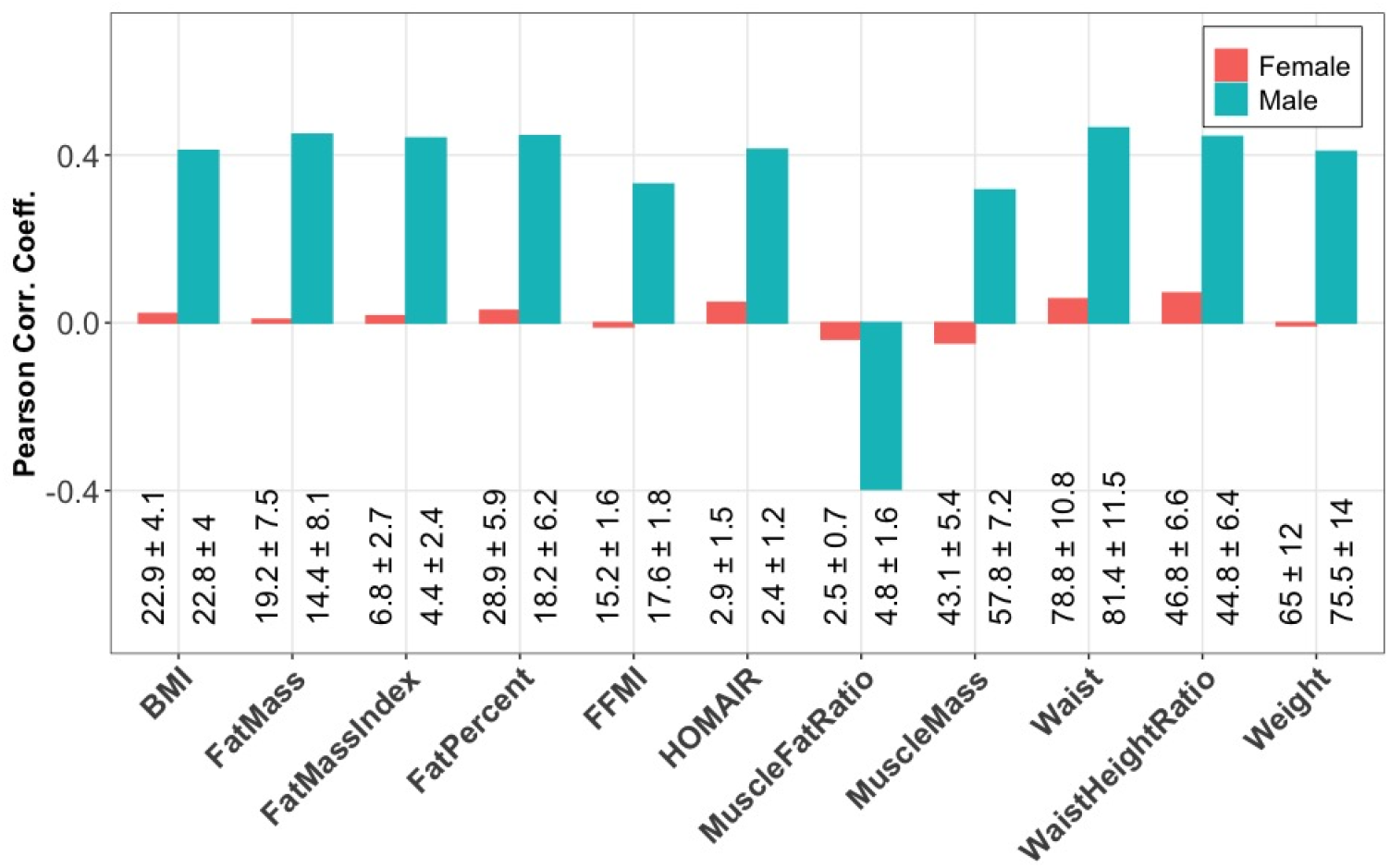
Correlation between the second component (**a**_2_) of the 2-component CP model of the T0-corrected data in the *subjects* mode with the additional meta data. Measurements from males and females are analyzed using a CP model separately as shown in Fig. 3 and Fig. 8. The second component vector in the *subjects* mode, i.e., (**a**_2_), from each model is used to compute the correlations for males and females. Descriptions of these variables are as follows: HOMAIR: Homeostatic model assessment for Insulin Resistance; MuscleFatRatio: Muscle to fat ratio; FatPercent: Body fat percentage; MuscleMass: Amount of muscle in the body (kg); Weight (kg); BMI: Body Mass Index; Waist: Waist circumferance (cm); WaistHeightRatio: Waist measurement divided by height (cm); FatMass: Amount of body fat (kg); FatMassIndex; FFMI: Fat Free Mass Index. The mean ± standard deviation for each variable and sex is shown at the bottom of the plot.

The first component, i.e., (**a**_1_, **b**_1_, **c**_1_), on the other hand, models an earlier response captured by **c**_1_ potentially modelling individual differences. Scatter plots of the first and second component in the *subjects* and *metabolites* modes showing subject groups and groups of lipoproteins based on their classes and subclasses are given in Fig. 6.

**Fig. 6.**
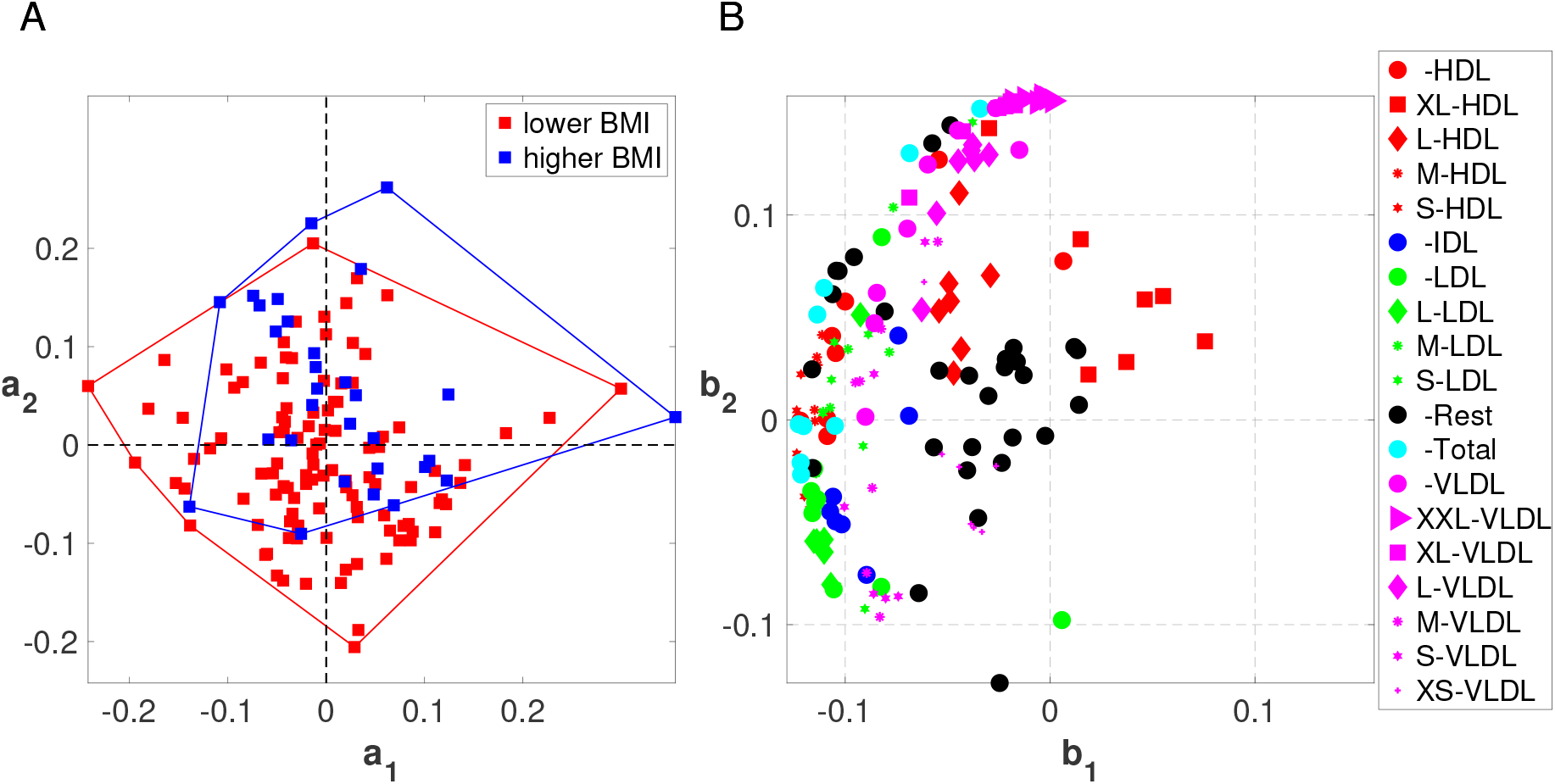
Scatter plots of the first and second component from the 2-component CP model of the T0-corrected data from males. (A) *Subjects* mode, (i.e., **a**_1_ and **a**_2_) colored according to BMI defined as lower BMI: BMI*<* 25 and higher BMI: BMI ≥ 25, and (B) *Metabolites* mode, where metabolites are colored according to lipoprotein classes. The size of the marker for each metabolite is adjusted according to lipoprotein subclasses as indicated in the legend. The Rest group contains features other than the ones in the lipoprotein group, the Total group corresponds to total concentrations of certain metabolites, e.g., Total-C, Total-TG. See supplementary material S1 for details.

### 3.2. Analysis of the fasting state metabolomics data from males using PCA reveals a BMI-related group difference

Fig. 7 shows the scatter plots of principal components in the *subjects* and *metabolites* modes from the PCA of the fasting state data. In the *subjects* mode, a weak but statistically significant (*p*-value = 1 × 10^*−*3^) group difference is captured using the first principal component (PC1). PC1 is plotted against the fourth PC (PC4) to facilitate the comparison with females (See subsection 3.4 for the fasting state analysis of females). In the *metabolites* mode, we observe that in PC1, IDLs, LDLs, VLDLs, Apolipoprotein B (ApoB), and Remnant cholesterol (Remnant-C) have high positive loadings showing positive relation with the concentrations of these metabolites and the higher BMI group.

**Fig. 7.**
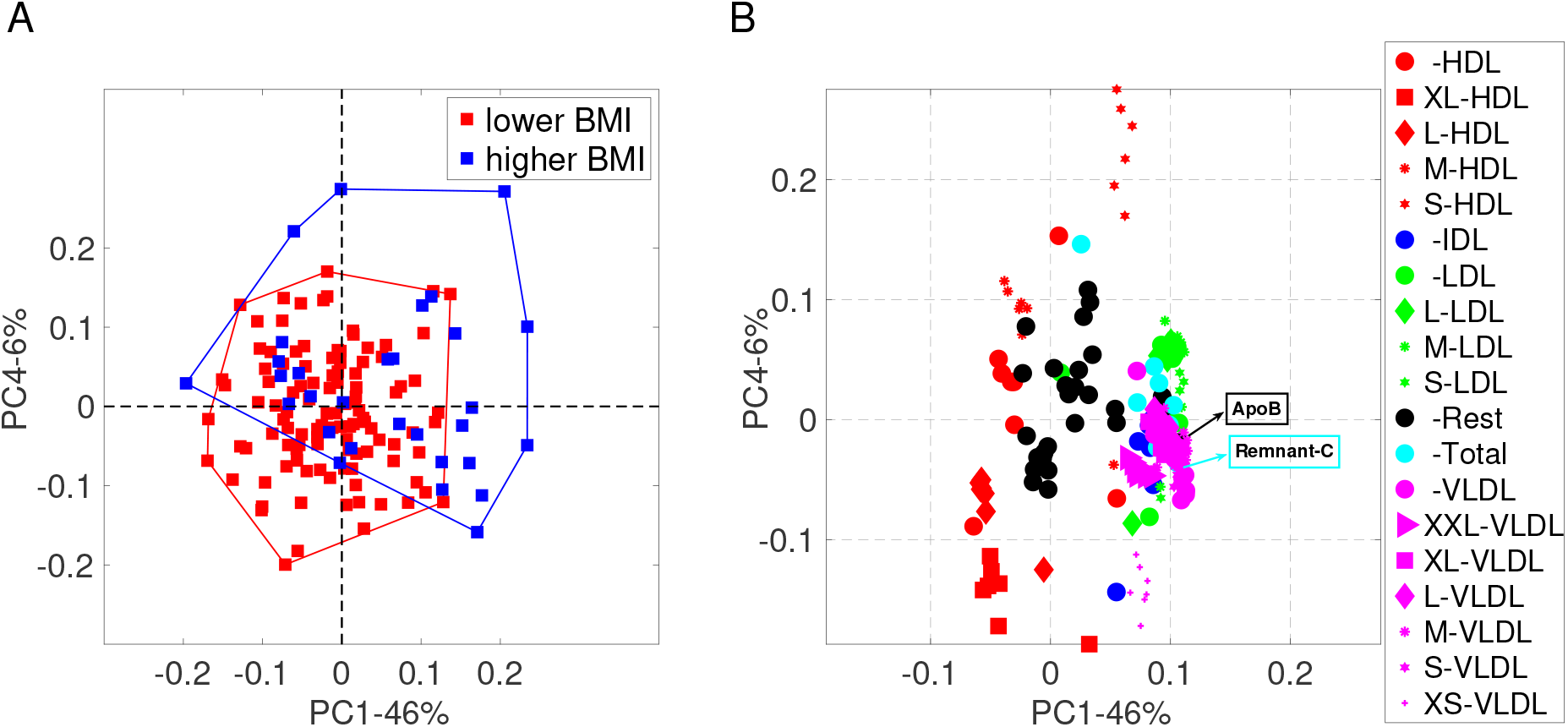
Scatter plots from PCA of the *fasting-state* data from males: (A) *Subjects* mode, where subjects are colored according to BMI defined as lower BMI: BMI*<* 25 and higher BMI: BMI ≥ 25, and (B) *Metabolites* mode, where metabolites are colored according to lipoprotein classes. The size of the marker for each metabolite is adjusted according to lipoprotein subclasses as indicated in the legend. We show the names of the metabolites with the highest coefficients for the first component, where we observe a statistically significant group difference in terms of BMI. The Rest group contains features other than the ones in the lipoprotein group, and the Total group corresponds to total concentrations of certain metabolites, e.g., Total-C, Total-TG. See supplementary material S1 for details.

### 3.3. The CP model of T0-corrected metabolomics data from females does not reveal a BMI-related group difference

Factors captured by the analysis of the T0-corrected data from females using a 2-component CP model are given in Fig. 8 (see supplementary material S2 for the selection of the number of components based on replicability). The fit of the 2-component model is 47%.

**Fig. 8.**
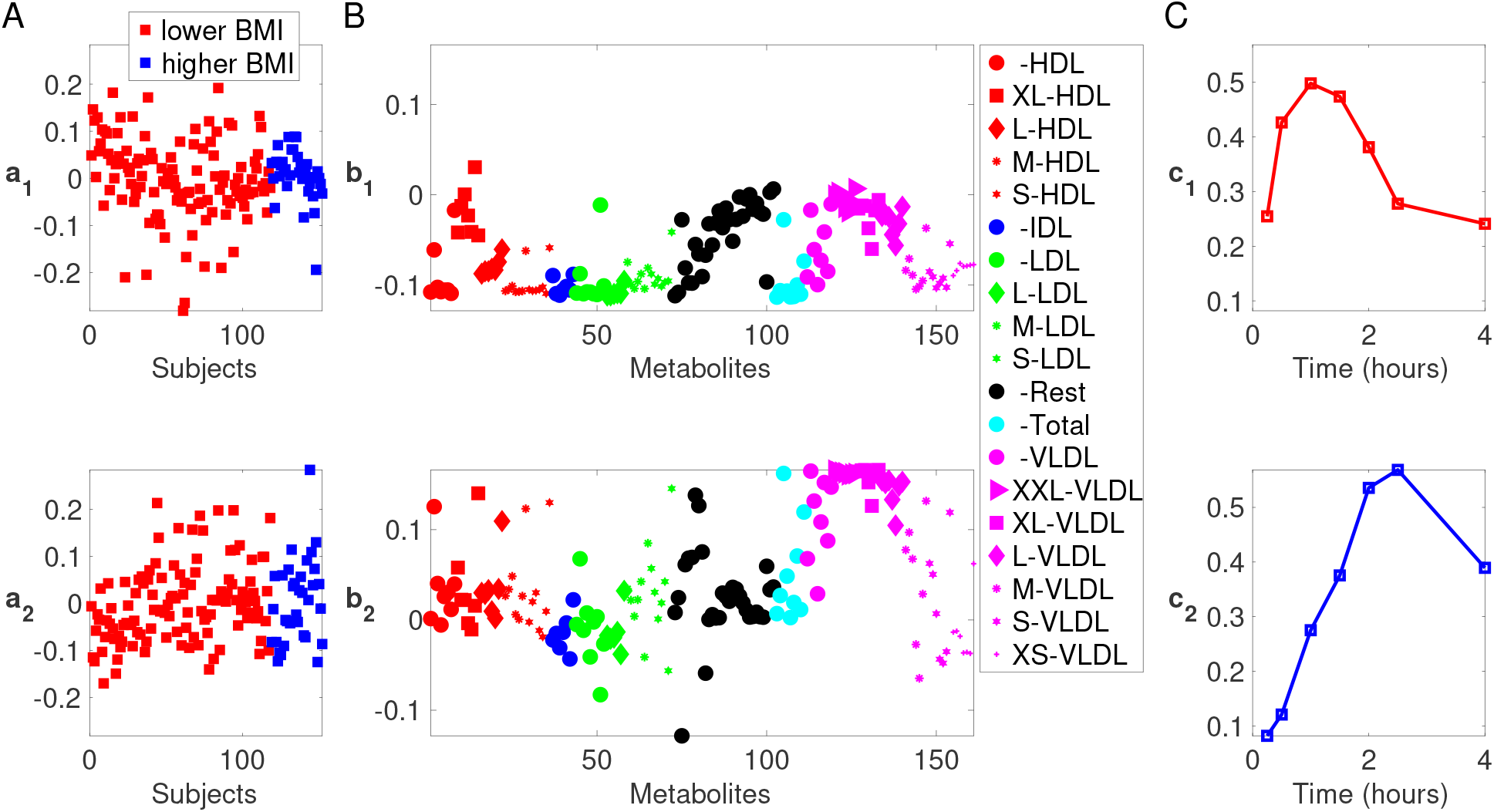
2-component CP model of the T0-corrected metabolomics data from females. (A) *Subjects* mode (i.e., **a**_1_ and **a**_2_), where subjects are colored according to BMI defined as lower BMI: BMI*<* 25 and higher BMI: BMI ≥ 25, (B) *Metabolites* mode (i.e., **b**_1_ and **b**_2_), where metabolites are colored according to lipoprotein classes. The size of the marker for each metabolite is adjusted according to lipoprotein subclasses as indicated in the legend. The Rest group contains features other than the ones in the lipoprotein group, the Total group corresponds to total concentrations of certain metabolites, e.g., Total-C, Total-TG, and (C) *Time* mode (i.e., **c**_1_ and **c**_2_).

In the *subjects* mode, we do not observe a group difference in either component in terms of the additional information available about the participants, see **a**_1_ and **a**_2_ in Fig. 8A, where subjects are colored according to *lower* and *higher* BMI groups. Note that correlations are quite low for females in Fig. 5. The *time* mode shows two underlying profiles: early response in **c**_1_, and a later response in **c**_2_. Even though the second component is similar in the *metabolites* and *time* modes, i.e., **b**_2_ and **c**_2_ in Fig. 8, to the component captured by the CP model of the T0-corrected data from males and revealing a BMI-related group difference, i.e., (**a**_2_, **b**_2_, **c**_2_) in Fig. 3, no BMI-related group difference is observed in this component for females (see also Fig. 10 for a clear comparison of the components from males vs. females).

**Fig. 9.**
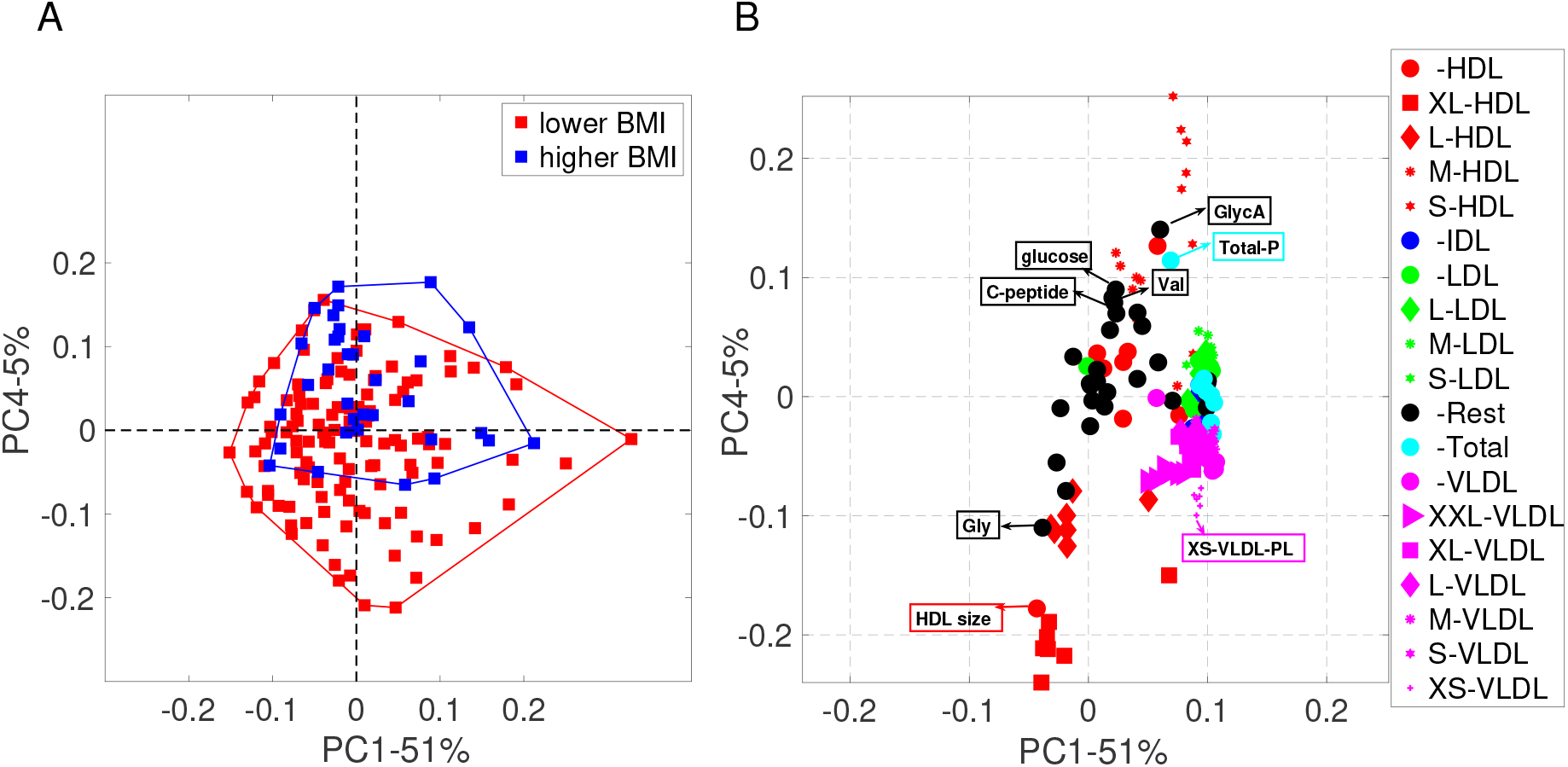
Scatter plots from PCA of the *fasting-state* data from females: (A) *Subjects* mode, where subjects are colored according to BMI defined as lower BMI: BMI*<* 25 and higher BMI: BMI ≥ 25, and (B) *Metabolites* mode, where metabolites are colored according to lipoprotein classes. The size of the marker for each metabolite is adjusted according to lipoprotein subclasses as indicated in the legend. We show the names of the metabolites with the highest coefficients for the fourth component, where we observe a statistically significant group difference in terms of BMI. The Rest group contains features other than the ones in the lipoprotein group, the Total group corresponds to total concentrations of certain metabolites, e.g., Total-C, Total-TG.

**Fig. 10.**
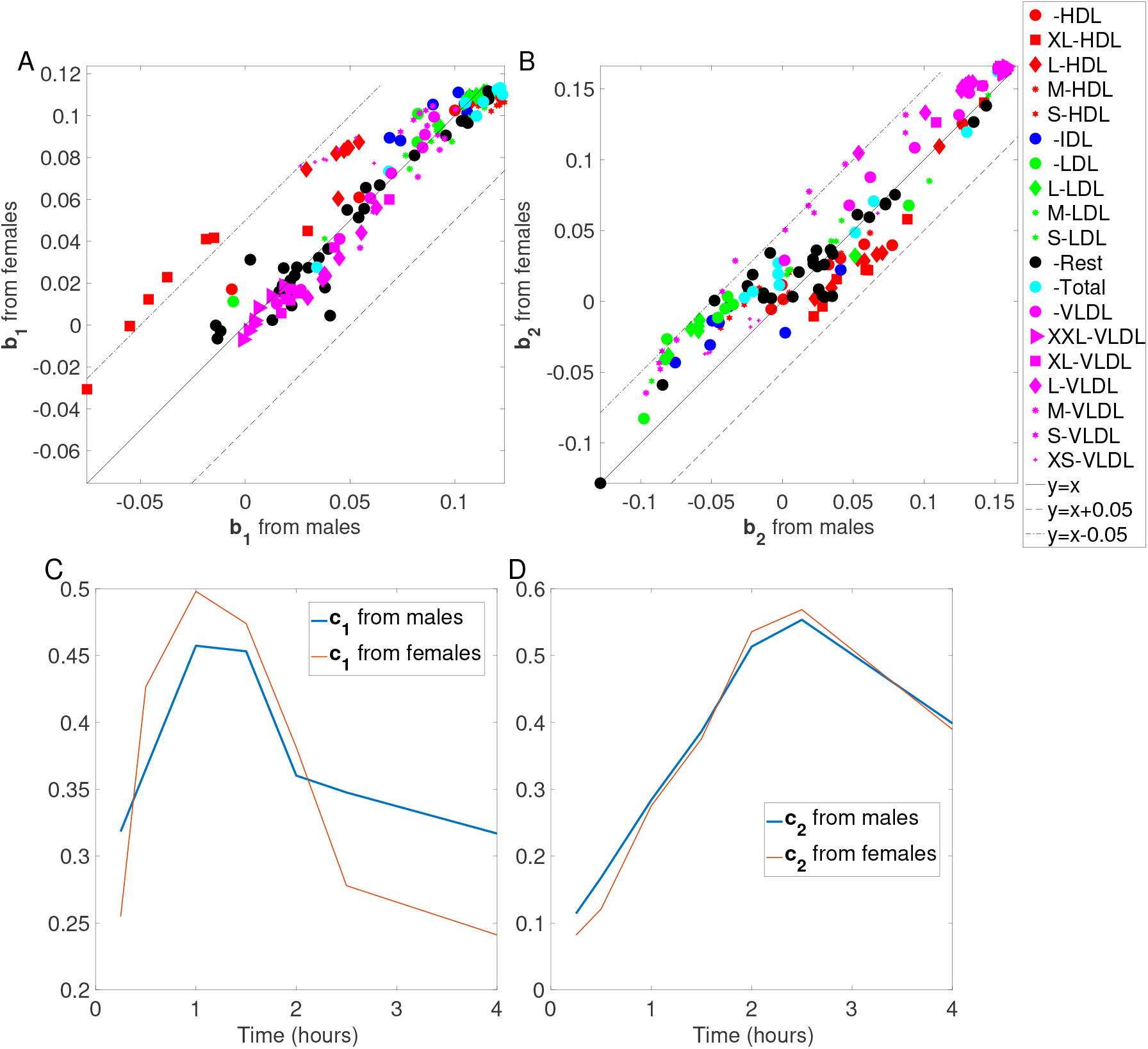
Comparison of CP models of the T0-corrected data from males and females. (A and B) *Metabolites* mode. To better illustrate the comparison between the patterns, the line *y* = *x* is plotted together with *y* = *x −* 0.05 and *y* = *x* + 0.05. (C and D) *Time* mode.

### 3.4. Analysis of the fasting state metabolomics data from females using PCA reveals a BMI-related group difference

Scatter plots of PC1 and PC4 in the *subjects* and *metabolites* modes from PCA of the fasting state data from females are shown in Fig. 9. In the *subjects* mode, only PC4 reveals a statistically significant (*p*-value = 5 × 10^*−*6^) BMI-related group difference. Note that the fourth component only explains 5% of the variation in the data so the signal - although statistically significant - is weak. In the *metabolites* mode, we observe that in PC4, HDLs (S), C-peptide, glucose, glycoprotein acetyls (GlycA), total concentration of lipoprotein particles (Total-P), and valine (Val) have high positive loadings (positively relate to *higher* BMI) while the HDLs (XL), average diameter for HDL particles (HDL size), glycine (Gly), and XS-VLDL-PL have high negative loadings (negatively relate to *higher* BMI).

We summarize the metabolites observed to be related to BMI-related group difference in Table 2.

**Table 2.** Metabolites with high coefficients (in terms of absolute value) observed to be related to BMI-related group difference in males vs. females. (+) and (-) denotes positively and negatively related to higher BMI.

|  | T0-corrected analysis | Fasting state analysis |
| --- | --- | --- |
| Males | (+): TG-related, VLDLs (XXL/XL/L), MUFA, SFA, and Total-FA<br>(-): LDL-FC-related, VLDLs (M/S), and unsaturation | (+): IDLs, LDLs, VLDLs, ApoB, and Remnant-C |
| Females |  | (+): HDLs (S), C-peptide, glucose, GlycA, Total-P, and Val<br>(-): HDLs (XL), Gly, HDL size, and XS-VLDL-PL. |

## 4. Discussion

### 4.A. Fasting vs. Dynamic states: In males, groups of lipoproteins reveal BMI-related group difference, and the relations of those lipoproteins with BMI groups differ in the fasting vs. dynamic state

In males, for the fasting state data, the positive relation of LDL-C (53, 54), VLDL-C (53) and ApoB (53) with higher BMI has been previously reported, and our findings are consistent with the literature. Likewise, we find similar patterns whereby large triglyceride rich VLDL lipoproteins are increased in individuals with higher BMI following a meal challenge containing fats assessed via conventional AUC postprandial metrics (55). However, we did not find complementary decreases in HDL, this may be in part due to gender stratification differences in our analyses. Likewise monozygotic twin studies point to that these observed differences in lipoprotein responses are driven by BMI differences, rather than underlying genetics (11).

While Table 2 gives a summary of the most important metabolites, i.e. metabolites with high coefficients in terms of absolute value, we also observe in Fig. 7 that HDLs (XL/L) negatively relate to higher BMI in the fasting state (even though they are not among the most important metabolites; therefore, not reported in Table 2). This matches with previous results in terms of HDL-C (53, 54).

This study differs from previous challenge test studies as it is a comprehensive study in terms of the lipoproteins (classes and subclasses) considered in the analysis and the compact summary provided by the CP model from the postprandial data in comparison with the fasting state analysis. We observe that dynamic and fasting state data provide different views of the same set of metabolites. These different views reveal different relations with the BMI groups as we observe in the case of VLDLs, i.e., while concentrations of all VLDLs positively relate to higher BMI in the fasting state (see Fig. 7), Fig. 3 shows that changes in VLDLs (XXL/XL/L) positively relate to higher BMI, and changes in VLDLs (M/S) negatively relate to higher BMI in the dynamic state.

These results not only improve our understanding of dynamic changes in the human metabolome but also reveal dynamic and static markers through the analysis of fasting and postprandial data.

### 4.B. Males vs. Females (Dynamic State): Patterns of dynamic response to the challenge test, i.e., patterns in the *metabolites* and *time* modes in T0-corrected analyses, are similar in males vs. females

Fig. 10 compares the components extracted using CP models from males vs. females in the *metabolites* and *time* modes. Fig. 10A and Fig. 10C show the comparison of **b**_1_s from males vs. females, and **c**_1_s from males vs. females, respectively. The second components (i.e., **b**_2_s and **c**_2_s), which are the patterns of interest in terms of BMI-related group difference, are compared in Fig. 10B and Fig. 10D. We observe similar patterns in both modes, especially for the second component. In the first component (Fig. 10A and Fig. 10C), slight differences are observed, e.g., HDLs (XL/L) and VLDLs (XS) in **b**_1_s are close to *y* = *x* + 0.05 suggesting slightly different metabolic responses in terms of these metabolites. Nevertheless, CP models of females and males show that patterns of dynamic response to the challenge test are similar in males and females.

We strengthen this argument by analyzing the T0-corrected data from both males and females simultaneously using a CP model. The 2-component CP model is given in supplementary material S3. When males and females are analyzed together using a CP model, the components in the *metabolites* and *time* modes are the same for both sexes. In the *subjects* mode, BMI-related group difference is observed among males but still not among females (See Figure S2 in supplementary material S3). The presence of these gender differences can be anticipated due to the inherent variances in sex hormones, psychosocial factors, and cardiovascular risk factor profiles (56). These postprandial gender differences may in part explain the cardiovascular risk reduction noted in pre-menopausal women (57).

### 4.C. Males vs. Females (Fasting State): In the fasting state, patterns in the *metabolites* mode are similar in males vs. females while the relations of those patterns with the BMI groups differ

Fig. 7 and Fig. 9 show the scatter plots of PC1 and PC4 from the fasting state analysis from males and females, respectively. For both males and females, similar patterns are observed. More precisely, in PC1s, IDLs, LDLs, and VLDLs are the metabolites with high positive coefficients while in PC4s the metabolites with high positive coefficients are mainly the HDLs (S) and with high negative coefficients are mainly the HDLs (XL), for both males and females. However, the BMI group difference is statistically significantly associated with PC1 in males while PC4 reveals a statistically significant group difference in terms of BMI in females.

That being said, for males, we also observe in PC1 (even though the coefficients are small) that the HDLs (S) positively relate to higher BMI while the HDLs (XL) negatively relate to the higher BMI group capturing the relation revealed by PC4 in females. Thus, for both males (from PC1) and females (from PC4), we can say that the HDLs (S) are positively related to higher BMI and HDLs (XL) are negatively related to higher BMI. However, we observe IDLs, VLDLs, LDLs playing a role in revealing group difference in males only (from PC1). These differences may be explained in the pre-menopausal cardioprotective lipid profiles of females (58). This gender disparity in postprandial atherogenic remnant lipoproteins may help explain why males are at greater cardiovascular risk in early life. Literature suggests gender differences in cardiovascular risk are abolished after menopause (59), and this may point towards the sex hormones playing a moderating role on atherogenic lipoproteins.

## 5. Conclusion

In this study, we have demonstrated a comprehensive analysis of metabolomics measurements collected during a challenge test from the COPSAC_2000_ cohort. We have arranged time-resolved metabolomics measurements as a *subjects* by *metabolites* by *time points* tensor, and analyzed fasting state-corrected dynamic data using a CP model in comparison with the fasting state data. Our analysis demonstrates that the CP model reveals interpretable patterns showing how subject groups (i.e., lower vs. higher BMI groups) differ in terms of the dynamic response of certain metabolite groups. We also show that metabolites behave differently in fasting vs. dynamic states; therefore, analysis of fasting and postprandial data can reveal static as well as dynamic biomarkers. In addition, our results indicate that patterns of dynamic response to the challenge test are similar in males vs. females but differences are observed in terms of how those patterns relate to BMI groups. Extracted patterns have shown to be not only interpretable but also replicable.

While in this paper we demonstrate that the CP model can reveal a compact summary of time-resolved postprandial data and reveal novel insights such as sex differences and metabolite groups behaving differently in fasting vs. dynamic states, there are still several limitations in terms of data analysis and findings. The set of measured metabolites/features is dominated by lipoproteins; therefore, the main patterns of variation in the data mainly model the behavior of lipoproteins. We do not observe glycolysis-related metabolites, amino acids or ketone bodies well-modelled by the CP model due to the dominant set of lipoproteins. In order to get a better understanding of postprandial metabolic response in terms of glycolysis, ketogenesis, and the aminoacid metabolism, metabolites may be grouped into specific subsets and jointly analyzed. We plan to explore this direction through the use of coupled tensor factorizations (60), which can jointly analyze multiple data sets in the form of higher-order tensors. Another potential limitation is that the CP model, as an unsupervised method, focuses on the main patterns that explain the variation in the data but these main patterns may not necessarily be related to meta variables of interest. Such an unsupervised approach may miss the relation with meta variables if that relation does not significantly contribute to the main variation in the data. In those cases, supervised methods may perform better. However, the goal of using an unsupervised approach in this exploratory study is its potential to reveal unknown subject stratifications rather than predicting certain meta variables. Finally, joint analysis of postprandial metabolomics data with other omics data sets, in particular, gut microbiome data, through data fusion methods (60, 61) is also of interest due to its promise to reveal better subject stratifications. With recent technological advances, e.g., multi-omics microsampling (62), that may make such challenge test data more easily available, jointly analyzing dynamic metabolomics data with other omics data sets becomes even more crucial.

## ACKNOWLEDGMENTS

We thank the children and families of the COPSAC_2000_ cohort for their contribution, and the clinical team at COPSAC for conducting the clinical testing.

## Footnotes

∗ The full list of metabolites is available at https://research.nightingalehealth.com/biomarkers.

