## Supplementary Materials for "Characterizing human postprandial metabolic response using multiway data analysis"

#### Supplementary Material S1

| | Cholesterol(C) | Triglycerides(TG) | Phospholipids(PL) | Cholesteryl esters (CE) | Free cholesterol (FC) | Total lipids(L) | Particles(P) | |
|---|---|---|---|---|---|---|---|---|
| Lipoproteins | VLDL | | | | | | | Size |
| | XXL-VLDL-C | XXL-VLDL-TG | XXL-VLDL-PL | XXL-VLDL-CE | XXL-VLDL-FC | XXL-VLDL-L | XXL-VLDL-P | XXL |
| | XL-VLDL-C | XL-VLDL-TG | XL-VLDL-PL | XL-VLDL-CE | XL-VLDL-FC | XL-VLDL-L | XL-VLDL-P | XL |
| | L-VLDL-C | L-VLDL-TG | L-VLDL-PL | L-VLDL-CE | L-VLDL-FC | L-VLDL-L | L-VLDL-P | L |
| | M-VLDL-C | M-VLDL-TG | M-VLDL-PL | M-VLDL-CE | M-VLDL-FC | M-VLDL-L | M-VLDL-P | M |
| | S-VLDL-C | S-VLDL-TG | S-VLDL-PL | S-VLDL-CE | S-VLDL-FC | S-VLDL-L | S-VLDL-P | S |
| | XS-VLDL-C | XS-VLDL-TG | XS-VLDL-PL | XS-VLDL-CE | XS-VLDL-FC | XS-VLDL-L | XS-VLDL-P | XS |
| | VLDL-C | VLDL-TG | VLDL-PL | VLDL-CE | VLDL-FC | VLDL-L | VLDL-P | |
| | Average diameter of VLDL | | | | | | | |
| | IDL | | | | | | | |
| | IDL-C | IDL-TG | IDL-PL | IDL-CE | IDL-FC | IDL-L | IDL-P | |
| | LDL | | | | | | | |
| | L-LDL-C | L-LDL-TG | L-LDL-PL | L-LDL-CE | L-LDL-FC | L-LDL-L | L-LDL-P | L |
| | M-LDL-C | M-LDL-TG | M-LDL-PL | M-LDL-CE | M-LDL-FC | M-LDL-L | M-LDL-P | M |
| | S-LDL-C | S-LDL-TG | S-LDL-PL | S-LDL-CE | S-LDL-FC | S-LDL-L | S-LDL-P | S |
| | LDL-C | LDL-TG | LDL-PL | LDL-CE | LDL-FC | LDL-L | LDL-P | |
| | Average diameter of LDL. | | | | | | | |
| | HDL | | | | | | | |
| | XL-HDL-C | XL-HDL-TG | XL-HDL-PL | XL-HDL-CE | XL-HDL-FC | XL-HDL-L | XL-HDL-P | XL |
| | L-HDL-C | L-HDL-TG | L-HDL-PL | L-HDL-CE | L-HDL-FC | L-HDL-L | L-HDL-P | L |
| | M-HDL-C | M-HDL-TG | M-HDL-PL | M-HDL-CE | M-HDL-FC | M-HDL-L | M-HDL-P | M |
| | S-HDL-C | S-HDL-TG | S-HDL-PL | S-HDL-CE | S-HDL-FC | S-HDL-L | S-HDL-P | S |
| | HDL-C | HDL-TG | HDL-PL | HDL-CE | HDL-FC | HDL-L | HDL-P | |
| | Average diameter of HDL | | | | | | | |
| | TOTAL | | | | | | | |
| | Total-C | Total-TG | Total-PL | Total-CE | Total-FC | Total-L | Total-P | |

| | |
|---|---|
| | Remnant-C and Total-FA |
| | **REST** |
| **Apolipoproteins** | ApoA1, ApoB |
| **Fatty acids** | Degree of unsaturation, Omega-3, Omega-6, DHA, LA, MUFA, PUFA, SFA |
| **Amino acids** | Ala, Gln, Gly, His, Ile, Leu, Total BCAA, Tyr, Val |
| **Glycolysis-related metabolites** | Citrate, Glucose, Lactate, Pyruvate |
| **Ketone bodies** | Acetate, Acetoacetate, Acetone, bOHbutyrate |
| **Other** | C-peptide, GlycA, Insulin |

### Supplementary Material S2
### Replicability of the CP model of the T0-corrected metabolomics data from females

Fig. S1 shows the replicability of the CP model of the T0-corrected data from females for different number of components, $R$. The model is replicable for $R = 1, 2, 4$. The 2-component CP model is chosen here because the 4-component model is degenerate, i.e., the Tucker's congruence (TC)[1] value given in Equation 1 between a pair of components is close to -1. For the 4-component CP model, the TC value is -0.78. The TC value between component $i$ and component $j$ is defined as follows:

$$\mathrm{TC}_{ij} = \frac{\mathbf{a}_i{}^\mathsf{T}\mathbf{a}_j}{\parallel \mathbf{a}_i \parallel \parallel \mathbf{a}_j \parallel} \frac{\mathbf{b}_i{}^\mathsf{T}\mathbf{b}_j}{\parallel \mathbf{b}_i \parallel \parallel \mathbf{b}_j \parallel} \frac{\mathbf{c}_i{}^\mathsf{T}\mathbf{c}_j}{\parallel \mathbf{c}_i \parallel \parallel \mathbf{c}_j \parallel}, \tag{1}$$

where $(\mathbf{a}_i,\ \mathbf{b}_i,\ \mathbf{c}_i)$ denotes the $i$th component of an $R$-component CP model.

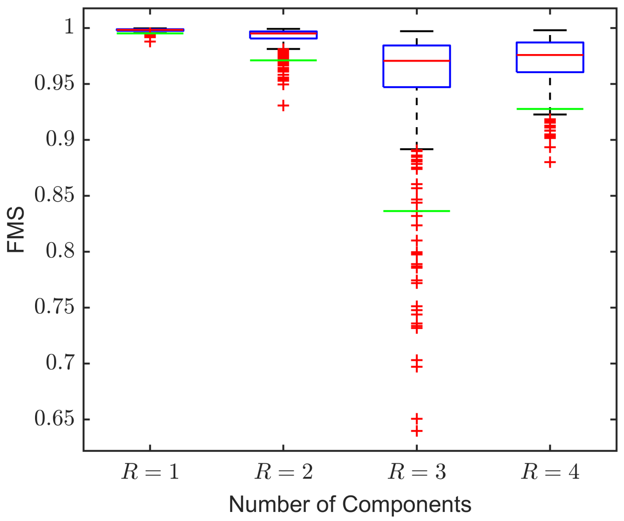


Figure S1: Replicability of the CP model of the T0-corrected data from females for different number of components, $R$. Green lines show that 95% of the FMS values are above that line. Models are replicable for $R = 1$, $R = 2$, and $R = 4$.

### Supplementary Material S3
### Comparison of CP models from Males vs. Females vs. All Subjects

#### 1 CP model of T0-corrected metabolomics data from all subjects

Fig. S1 shows the 2-component CP model of the T0-corrected metabolomics data from all subjects, i.e., males and females. The model fit is 45%.

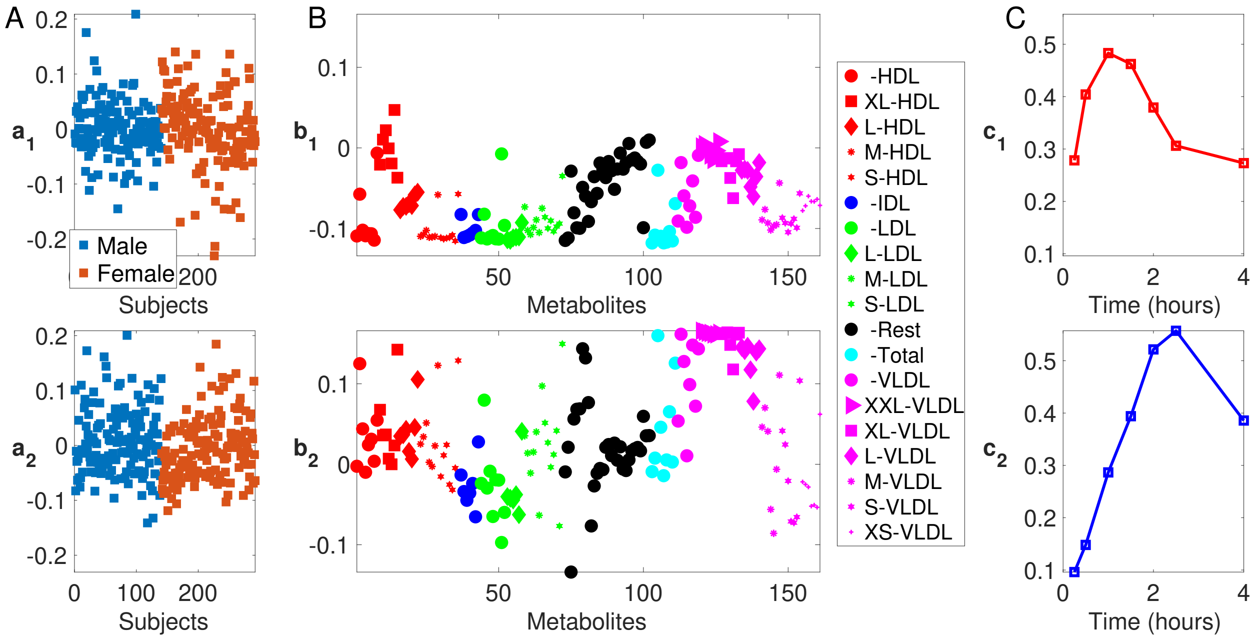


Figure S1: 2-component CP model of the T0-corrected metabolomics data from all subjects. (A) *Subjects* mode (i.e., $\mathbf{a}_1$ and $\mathbf{a}_2$). (B) *Metabolites* mode (i.e., $\mathbf{b}_1$ and $\mathbf{b}_2$). (C) *Time* mode (i.e., $\mathbf{c}_1$ and $\mathbf{c}_2$).

The CP model does not reveal any gender-related group difference. The components in the *metabolites* and *time* modes are similar to the components extracted using CP models from only males and only females. This supports our finding that males and females have similar patterns of dynamic response to the challenge test. In the second component, which is similar to the second component of CP models from only males and only females, there is a BMI-related group difference among males (with a $p$-value $= 6 \times 10^{-4}$) but not among females. See the boxplots (using $\mathbf{a}_2$) in Fig. S2.

#### 2 Patterns of dynamic response to the challenge test are similar in males vs. females vs. all subjects.

Here, we compare the components extracted from the *metabolites* and *time* modes of T0-corrected data using CP models from males vs. females vs. all subjects. Fig. S3 shows $\mathbf{b}_1$ and $\mathbf{b}_2$ from the three CP models. We observe that the components from these three CP models are very similar, except the metabolites mentioned in Section 4.B. in the manuscript. Fig. S4 demonstrates $\mathbf{c}_1$ and $\mathbf{c}_2$ from the three CP models showing that similar temporal patterns are extracted from all data sets.

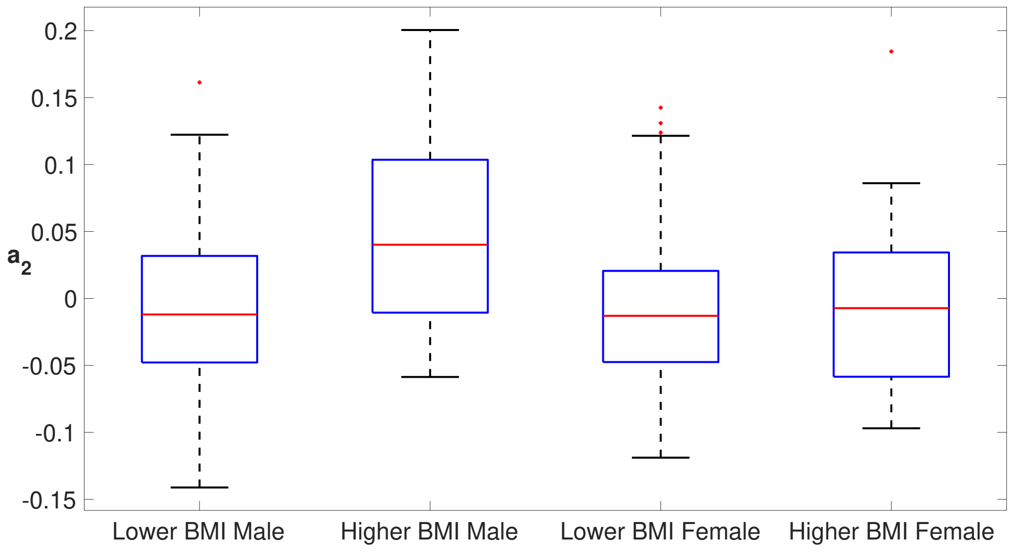


Figure S2: Boxplots of $\mathbf{a}_2$ from the 2-component CP model of the T0-corrected data from all subjects.

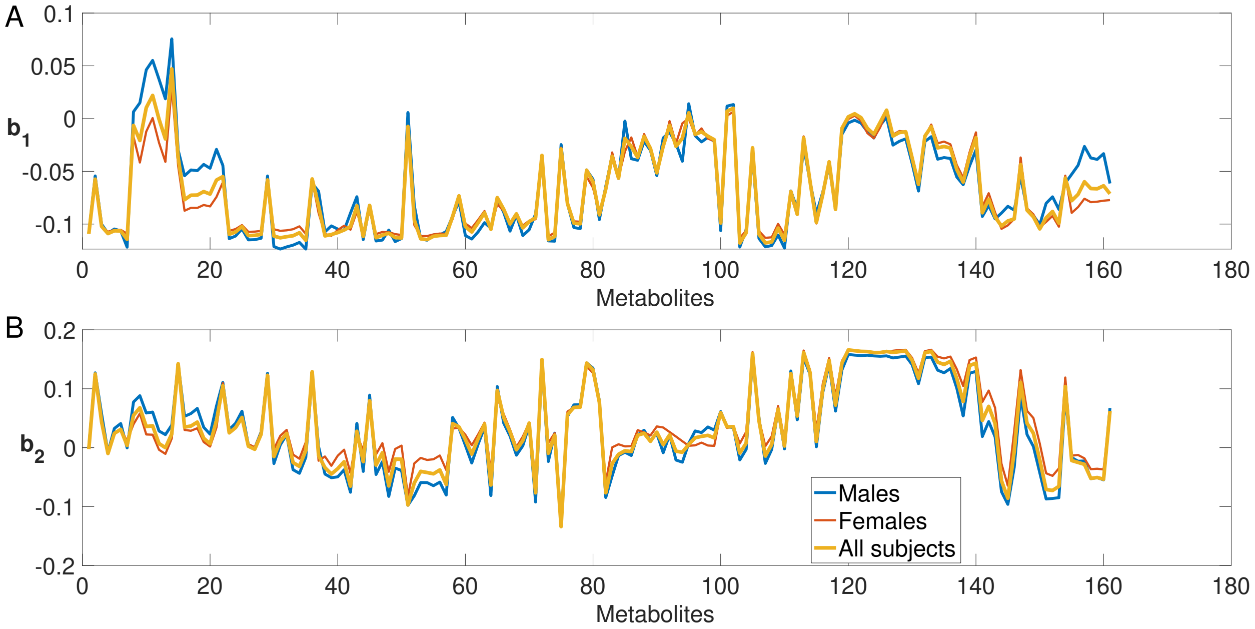


Figure S3: Comparisons of the *metabolites* modes using the three CP models from males vs. females vs. all subjects. (A) $\mathbf{b}_1$, (B) $\mathbf{b}_2$.

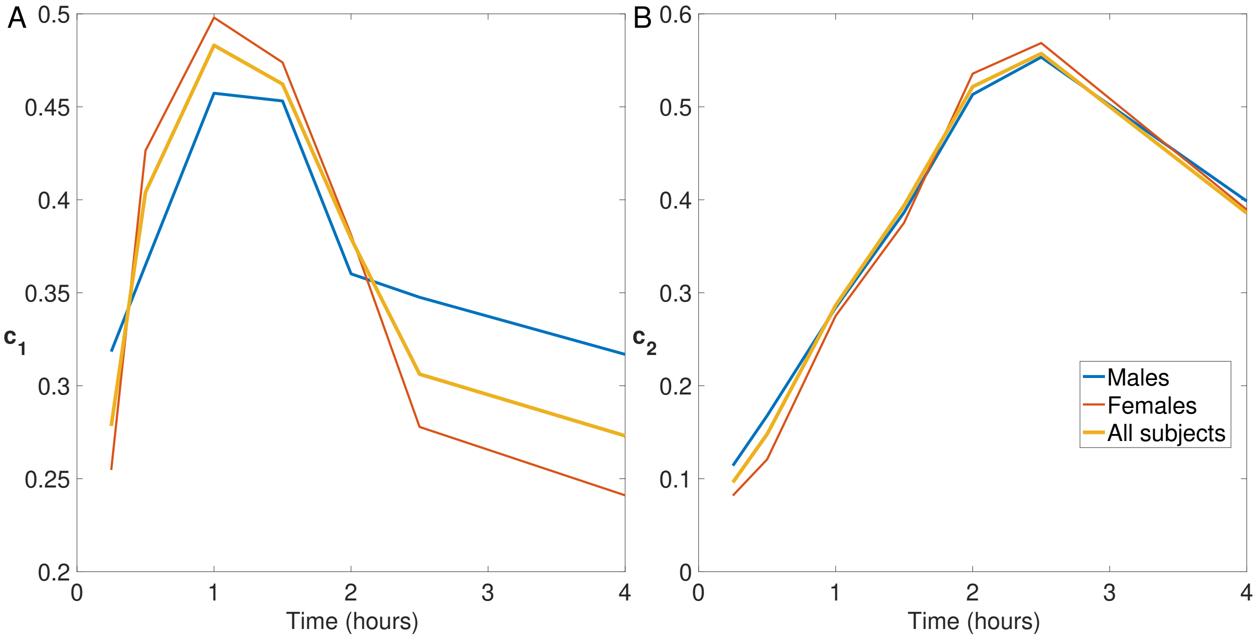


Figure S4: Comparisons of the *time* modes using the three CP models from males vs. females vs. all subjects. (A) $\mathbf{c}_1$, (B) $\mathbf{c}_2$.
